# Accurate Name Entity Recognition for Biomedical Literatures: A Combined High-quality Manual Annotation and Deep-learning Natural Language Processing Study

**DOI:** 10.1101/2021.09.15.460567

**Authors:** Dao-Ling Huang, Quanlei Zeng, Yun Xiong, Shuixia Liu, Chaoqun Pang, Menglei Xia, Ting Fang, Yanli Ma, Cuicui Qiang, Yi Zhang, Yu Zhang, Hong Li, Yuying Yuan

## Abstract

A combined high-quality manual annotation and deep-learning natural language processing study is reported to make accurate name entity recognition (NER) for biomedical literatures. A home-made version of entity annotation guidelines on biomedical literatures was constructed. Our manual annotations have an overall over 92% consistency for all the four entity types — gene, variant, disease and species —with the same publicly available annotated corpora from other experts previously. A total of 400 full biomedical articles from PubMed are annotated based on our home-made entity annotation guidelines. Both a BERT-based large model and a DistilBERT-based simplified model were constructed, trained and optimized for offline and online inference, respectively. The F1-scores of NER of gene, variant, disease and species for the BERT-based model are 97.28%, 93.52%, 92.54% and 95.76%, respectively, while those for the DistilBERT-based model are 95.14%, 86.26%, 91.37% and 89.92%, respectively. The F1 scores of the DistilBERT-based NER model retains 97.8%, 92.2%, 98.7% and 93.9% of those of BERT-based NER for gene, variant, disease and species, respectively. Moreover, the performance for both our BERT-based NER model and DistilBERT-based NER model outperforms that of the state-of-art model—BioBERT, indicating the significance to train an NER model on biomedical-domain literatures jointly with high-quality annotated datasets.

## Introduction

With the rapid development of next-generation sequencing technology, the cost of interpreting the clinical significance of hundreds of thousands of genomic variants has become an obvious bottleneck for the whole genetic testing process.^1-3^ There are dozens of well-established biological databases that are curated and maintained by researchers, which facilitate the interpretation of genomic variants by clinicians, geneticists and biologists. However, the information provided by these valuable data resources is still quite limited.^4-9^ Literatures in biomedical domain are instead a huge repository to store tremendous knowledge for genetic variant interpretation and they are continuously updating. It indeed poses great challenge for genetic interpreters to consult literatures manually to find relevant literature evidences for a new variant. To our best knowledge, literature evidence search for an unknown variant is a rate-determining step for genetic variant interpretation. Therefore, it should be quite helpful to make literature evidence searching more efficient, not to mention to fully automate the searching task. In the biomedical domain, one primary application of natural language processing (NLP) is to identify key concepts in literatures,^10-13^ which is the first step to both filter relevant articles based on contents for literature evidence searching and develop an automated literature evidence searching tool.^14^

As the main tool of identifying concepts in free texts, name entity recognition (NER) has long received a good deal of attention in NLP.^10,15-18^ Web-based services such as Pubtator, LitVar and Pubtator Central (PTC) were launched subsequently to automate annotations of literatures by combining existing text mining tools using rule-based and machine-learning-based NER techniques.^4,19-22^ The F1-scores for gene, variant, disease and species that PTC achieves were reported as 86.70%, 86.24%, 83.70% and 85.42%, respectively.^19^ It is obvious that there is a lot of room to improve the performance of such techniques due to their limitation of contextualized information.^23^ In the past few years, deep learning neural network (DNN) such as bidirectional long and short term memory (BiLSTM) combined with conditional random field (CRF) have greatly improved performance in NER, but the constraints of sequential computations remain a problem.^11-13,24-26^ In 2018, Google proposed a new self-attention-based language representation model called BERT, which pretrains deep bidirectional representations from unannotated texts and then finetunes them on annotated texts to create state-of-the-art models for a wide range of NLP tasks.^27^ In 2019, BioBERT was reported to pretrain and finetune pretrained BERT representations on biomedical texts, demonstrating that it is crucial to pretrain BERT on biomedical corpora when applying it to the biomedical domain.^28^ However, due to the lack of a high-quality dataset with all interested entity types annotated on the same corpus, BioBERT was trained for each entity type separately. Moreover, BioBERT was not applied to recognize variant for the shortage of the variant-annotated corpus, although variant is an extremely important entity type in genetic variant interpretation.^28^ In the same year, BioBERT team also introduced the web-based tool called BERN to tag entities in PubMed articles or raw texts, relying on tmVar 2.0 to extract variant while BioBERT for all the rest. ^29^

In spite of the high performance of BERT-based NER models, such large and complicated models face serious challenges when it comes to on-device real-time applications or under constrained computational training or inference budgets.^30^ A key solution to this problem in artificial intelligence (AI) community is knowledge distillation, in which a small model - the student - is trained to keep the same knowledge of a larger model - the teacher.^30^ There are a bunch of distilled versions of BERT such as BERT-PKD,^31^ DistilBERT,^32^ TinyBERT,^33^ and BERT-EMD.^34^ It was reported that DistilBERT reduces the size of a BERT model by 40%, while retaining 97% of its language understanding capabilities and being 60% faster. ^32^

Here we report a combined high-quality manual annotation and deep-learning NLP study to make accurate NER for biomedical literatures. A home-made version of entity annotation guidelines on biomedical literatures was constructed. The interested entity types include gene, variant, disease and species, which are all critical for genetic variant interpretation. The performance of our annotation was assessed by comparing our annotated results with those publicly available from experts previously.^20,21,35-38^ The final consistency for all the four entity types is over 92%. The possible reasons for the 8% discrepancy are also discussed. A total of 400 full biomedical articles from PubMed are annotated based on our home-made entity annotation guidelines. All the manually annotated corpora as well as approximately 500,000 BERN-annotated PubMed abstracts are used as a critical part of the datasets for deep learning NER model development. Both a BERT-based large model and a DistilBERT-based simplified model were constructed, trained and optimized for offline and online inference, respectively. Offline inference refers to an approach that ingests all the data at one time to do model prediction whereas online inference is the one that ingests data one observation at a time. Offline inference can be potentially applied to knowledge base construction while online inference can be used in to build interactive prediction tools. The F1-scores of the DistilBERT-based NER model retains 97.8%, 92.2%, 98.7% and 93.9% of those of BERT-based NER for gene, variant, disease and species, respectively. Moreover, the performance for both our BERT-based NER model and DistilBERT-based NER model outperforms that of BioBERT, indicating the significance to train an NER model on biomedical literatures jointly with high-quality annotated datasets.

## Methods

### Annotated Data Acquisition

Annotated literatures in this study were obtained in two ways: (1) our manual annotation, (2) downloading from public resources. The details are as follows:

#### 1, Our manual annotation

Our annotation was focused on four entity types, namely, gene, variant, disease and species. The annotating procedure was based on our home-made entity annotation guidelines on biomedical literatures, which was developed by four annotators in our group. These annotators all have had at least five-year experience in interpreting genetic testing reports at Beijing Genomics Institute (BGI). The overall procedure of the development of our home-made entity annotation guidelines can be described here briefly. Firstly, 10 common articles were assigned to each of the four annotators to annotate independently based on their own knowledge and public databases such as HGNC, NCBI. Taxonomy, Mondo Disease Ontology, Orphanet, etc. The four annotators then compared, discussed and analyzed their annotated results to form a preliminary version of the annotation guidelines. Subsequently, such steps were iterated in two additional rounds to finalize the annotation guidelines. In order to validate our home-made version of the annotation guidelines, our annotators finally annotated part of four publicly available corpora that were annotated by experts previously for each of the four entity types^20,21,36,37^ and made the consistency analysis.

Based on the above home-made entity annotation guidelines, a total of 400 full biomedical articles on PubMed were annotated. The overall workflow of manual annotation is shown in Figure 1. Briefly, our interested literatures were searched on PubMed and annotated with PTC,^19^ serving as the initial corpus for manual annotation. In the early stage of our annotation, the strategy of three-person manual annotation was adopted since annotators were not quite familiar with the whole annotation guidelines. Specifically, two annotators were assigned the same batch of articles independently. If their annotations of a complete article agree with each other, the annotated article would be ready for random inspection; otherwise, their discrepant annotations would go to a reviewer for correction before being added to the batch of articles for random inspection. It is noted that the reviewer also gave the feedback to the original annotators to make sure their annotation strategies would be well aligned over time. In the middle and late stages of our annotation, the strategy of two-person manual annotation is employed for a higher efficiency, in which only one annotator and one reviewer were involved for annotation. As for random inspection, a certain number of annotated articles in a batch were random inspected. If any problem arose, the whole batch would go back to the reviewer for recheck; otherwise, the whole batch would be finally aggregated into the annotated corpus when quality control is completed.

**Figure 1.**
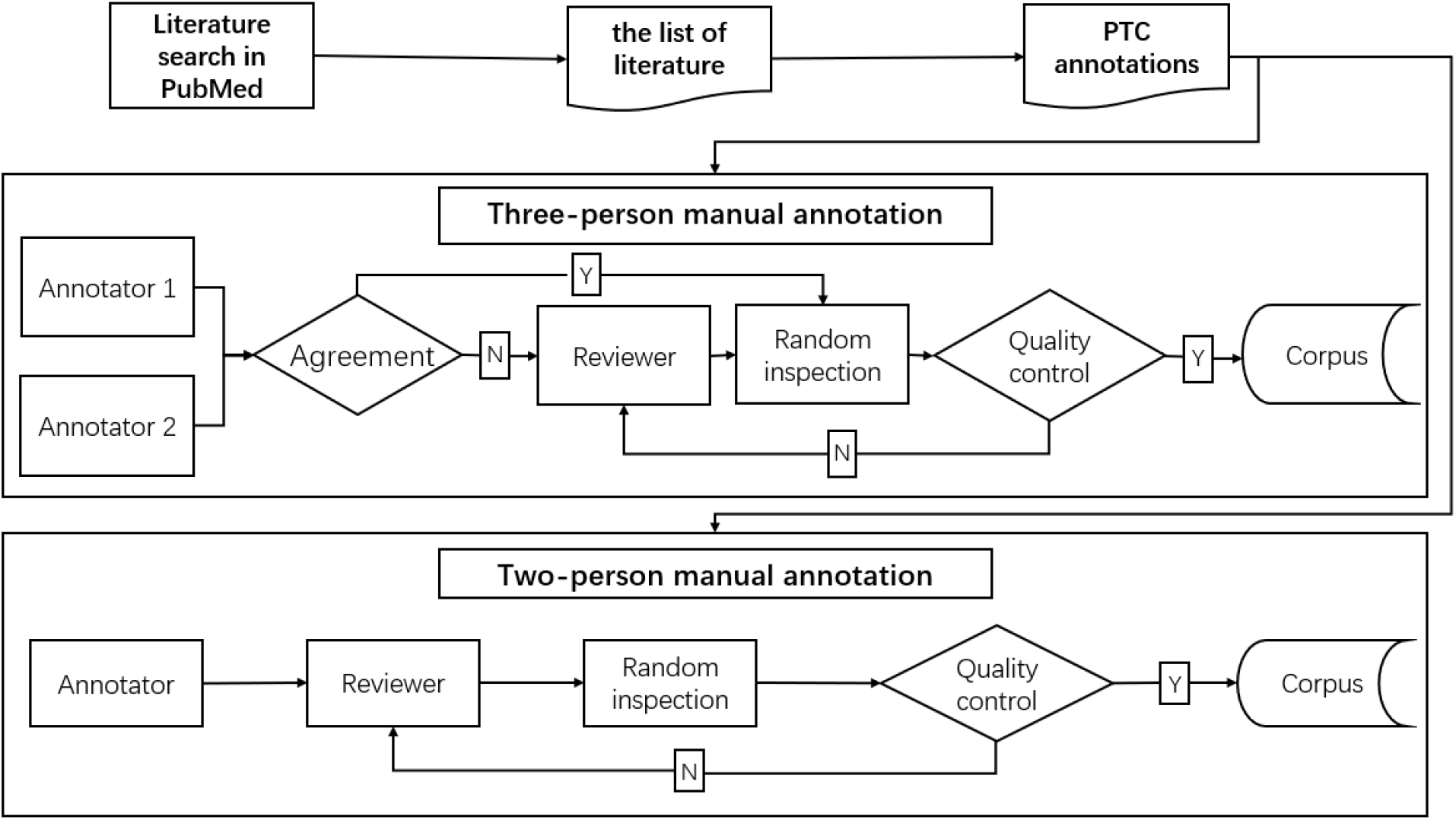
the workflow of manual annotation

Generally speaking, it is a time-consuming process to annotate literatures manually. In our experience, it took 4 annotators approximately 120 hours to complete the annotation of all the 400 biomedical literatures.

#### 2, Downloading from public resources

Approximately 500,000 BERN-annotated PubMed abstracts were downloaded via https://bern.korea.ac.kr/.

### Data preprocessing

Approximately 500,000 BERN-annotated PubMed abstracts were selected as datasets in this study, corresponding to the first 18 folders of the total of 1200 downloaded ones. Due to the sparsity of the overall annotated variants in the downloaded PubMed abstracts, the sentences containing the entity type of variant from all 1200 folders were collected. Both our annotated 400 articles and BERN-annotated abstracts were divided into train/validation/test datasets at the ratio of 7:2:1 for model training, validating and testing. All the corpora were converted into CoNLL format and labeled using BIO format.^39^

### NER Model Building

In order to build NER predictive models for both scenarios such as offline inference and online inference, a large model and a compact model were designed correspondingly, namely, BERT-based NER and DistilBERT-based NER, as is show in Figure 2. The overall process of the NER models contains two stages: pre-training and fine-tuning with two phases, as is illustrated in Figure 2(a).

**Figure 2.**
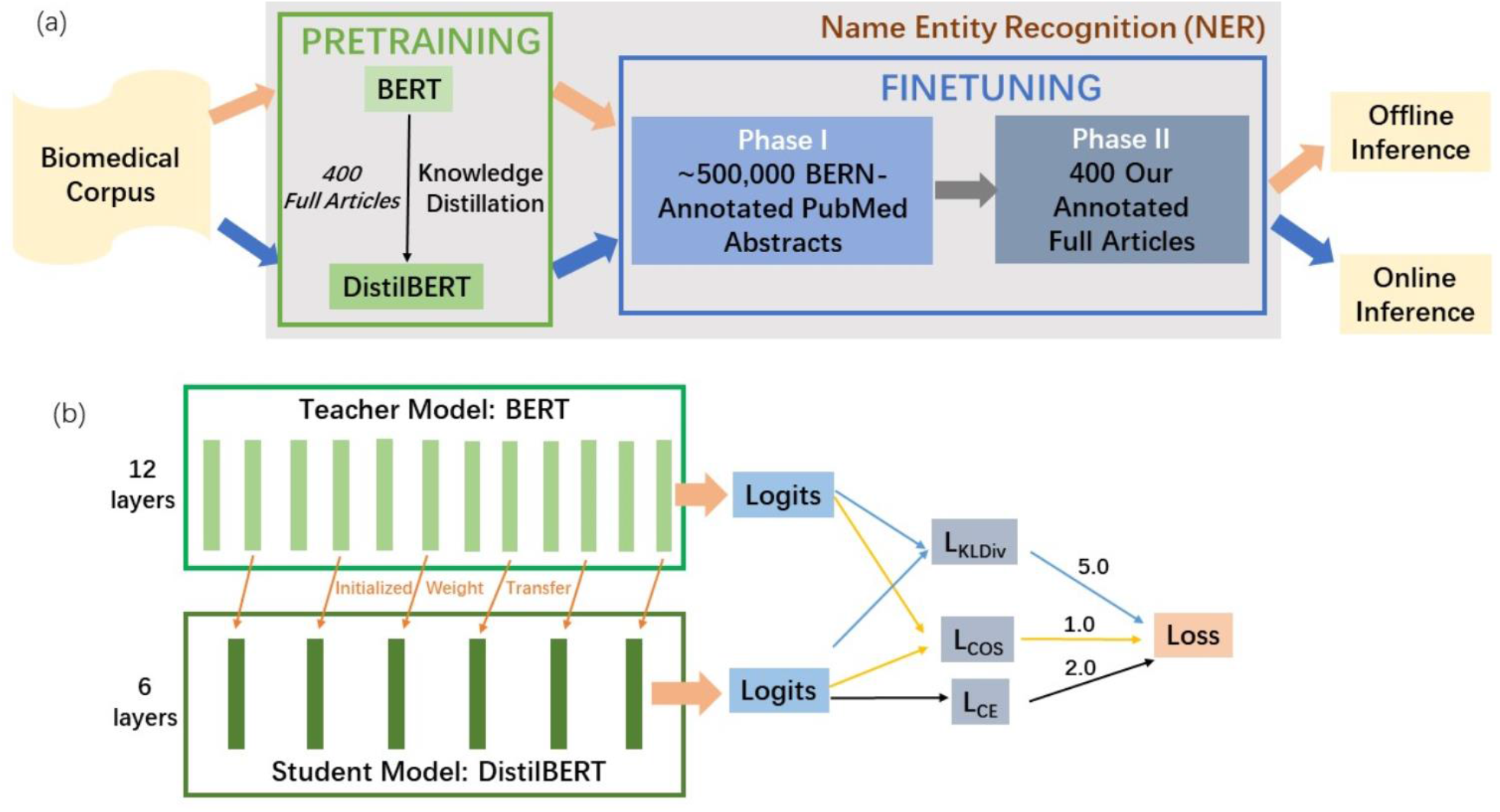
(a) the schematic of BERT-based and DistilBERT-based name entity recognition (NER) strategies, corresponding to the orange and blue arrows, respectively. In general, the BERT-based model is much larger than the DistilBERT-based model so that the former better fits offline inference while the latter can serve online inference. The NER module contains the pretraining stage and the finetuning stage with two phases due to the relatively small data set size for the second finetune phase. The pretrained weights of BERT NER model are from Biobert while those of DistilBERT are distilled from Biobert using 400 full articles, which is also used for Phase II at the finetuning stage. (b) The structural details of knowledge distillation. the teacher model (BERT) contains 12 layers while the student model (DistilBERT) has 6 layers. The well pretrained weights of the 2nd, 4th, 6th, 8th, 10th and 12th layers of the teacher model are transferred as the initialized weight of the student model. The output logits of the last layers of both the teacher and student models are used to calculate the total loss of the model according to *Loss*=5.0*\*L*_*KLDiv*_ +1.0 *\*L*_*COS*_ +2.0 *\* L*_*CE*._

#### 1, BERT-based NER

The BERT-based NER model in this study is a pretrained language representation model based on BERT for biomedical literatures. At the pre-training stage, the BERT-based NER model was loaded with weights from BioBERT, which was pretrained on 1 million PubMed abstracts based on BERT. The finetuning stage contains two phases with Phase I corresponding to the finetuning process of ∼500,000 BERN-annotated abstracts and Phase II corresponding to finetuning our 400 annotated full articles. The optimized BERT-based NER model is a large deep learning model with approximately 110 M parameters and expected to take relatively long time to run.

Similar to BioBERT, wordpiece embeddings that divide a word into several sub-words were employed in BERT-based NER model so as to recognize both known and out-of-vocabulary entities. The overall architecture of the BERT-based model is the same with BioBERT characteristic by 12 encoder layers and 12 attention heads. Different encoder layers and attention heads have been proven to capture different levels of input features such as surface, syntactic and semantic information.^27^ There are three major modifications of the current BERT-based NER model compared to BioBERT. Firstly, instead of only one phase in the finetuning stage, the current BERT-based NER model has two phases for finetuning. That is, the pretrained weights from BioBERT were first finetuned on ∼500,000 BERN-annotated PubMed abstracts so as to achieve roughly acceptable weights before being further finetuned on the 400 annotated full articles. Secondly, the labels for the current BERT-based NER model include the entity type of variant, which is extremely significant to understand genetic diseases but missing in BioBERT due to the shortage of a high-quality annotated dataset. Finally, unlike separate prediction of each entity type in BioBERT, the current BERT-based NER model trained all four annotated entity types in a single model, generating representations that capture invariant properties to tasks by sharing features.

#### 2, DistilBERT-based NER

The strategy of knowledge distillation in this study is through a general-purpose pre-training distillation. In one word, the student model DistilBERT obtains pretrained weights from the distillation of that of the teacher model BERT at the pre-training stage while maintaining the finetune stage the same as that of BERT-Based NER model. As a distilled version of BERT, DistilBERT is characteristic by the overall same architecture as BERT with only half the number of its layers while the token-type embeddings and the pooler are removed.^32^ As is seen from Figure 2(b), the architecture of the teacher BERT has 12 encoder layers, and thus the student DistilBERT has 6 encoder layers. The student model is initialized from the teacher model by taking the latter one layer out of two and is trained to reproduce the behavior of teacher model. The training loss is given by^32^

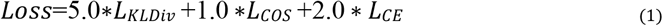

Where *L*_*kLDiv*_ is Kullback-Leibler divergence loss between the soft target probabilities of the teacher and the student, *L*_*COS*_ is the cosine embedding loss between the soft target probabilities of the teacher and the student and *L*_*CE*_ is the cross-entropy loss of the soft target probabilities of the student.

### NER Model Evaluation

To evaluate the performance of BERT-based and DistilBert-based NER models, reported metrics of BERN were compared and BERN was also applied to the same test set in the current study. It is noted that the NER of BERN for gene, disease and species relies on BioBERT while variant extraction adopts tmVar 2.0.^29^ The tool can be accessed at https://github.com/dmis-lab/bern. As for the evaluation metrics of NER, entity-level precision, recall and F1 score were calculated.

## Results

### 1, Consistency analysis between our annotators and experts annotating the public available corpus

Table 1 displays the statistics of the annotations from our annotators, publicly available annotated corpora from experts previously, the intersection between these two parties and the consistency rate. In total, 3,818 genes, variants, diseases and species were annotated from us while the total number of the annotations from experts previously is 3,868, resulting in an overall consistency rate of 92.24%. Specifically, the consistency rates for gene, variant and specie are all over 94.00%, among which those of gene and species are both as high as around 98% while that for disease is only 76.59%. The list of all the inconsistent cases are provided in supplementary materials.

**Table 1.** the statistics of the annotations from our annotators, publicly available annotated corpora from experts previously, the intersection between these two parties and the consistency rate.

| Entity Type | Dataset | Number of Annotations |  |  | Consistency Rate (%)<br>(#Intersection/<br>#Experts) |
| --- | --- | --- | --- | --- | --- |
|  |  | Our annotators | Experts | Intersection |  |
| Gene | GnormPlus/BioCreative II GN | 1,269 | 1,256 | 1,232 | 98.09 |
| Variant | tmVar 2.0 | 509 | 464 | 437 | 94.18 |
| Disease | NCBI disease | 844 | 961 | 736 | 76.59 |
| Species | Linnaeus | 1,196 | 1,187 | 1,163 | 97.98 |
| Total |  | 3,818 | 3,868 | 3,568 | 92.24 |

Table 2 shows the statistics of inconsistent annotated entities between experts previously and our annotators due to three different factors (discrepant rules of both annotation parties, the false annotations from the experts and our false annotations), the total inconsistent number, experts’ false annotation rate and the discrepant rules rate. The most significant factor that causes annotation inconsistency between experts and us for each entity is bolded. It is obvious that discrepant rules between previous experts and our annotators should be the dominant factor for all the inconsistent annotated entities and the ratio of inconsistent cases caused by discrepant rules to total inconsistent cases for gene, variant, disease and species is 58.33%, 96.30%, 79.17% and 71.11%, respectively. For instance, a mention without a clear description of a specific variant was annotated as variant by previous experts while not annotated by our annotators such as “T>C” (PMID:20738799), “G/A” (PMID:20738799), “G>C” (PMID:17961316) and “C-->T” (PMID:16001362). Detailed information for all inconsistent cases is provided in supplementary materials. In addition, false annotating from experts is the second most prominent factor for the inconsistency of disease annotation, which contributes to 25.78% annotation inconsistency. According to our annotation guidelines, an entity that either refers to a major category of disease, or is not curated in our referenced databases or is not curated as a phenotype should not be annotated as the entity type of disease. However, such entities are instead annotated as disease by previous experts. For example, “cancer” was not annotated as the entity type of disease from us while it was annotated in NCBI disease corpus. In addition, the inconsistency caused by the false annotation from experts is also as high as 25.78%. The main reason is that the entities that should belong to the entity type of gene are annotated as disease. For example, “APC” should be a gene in semantics while it was actually annotated as disease in NCBI disease (PMID:9724771).

**Table 2.** the statistics of inconsistent annotated entities between experts previously and our annotators due to three different factors (discrepant rules of both annotation parties, the false annotation from the experts and our false annotation), the total inconsistent number, experts’ false annotation rate and the discrepant rules rate. The most significant factor that causes annotation inconsistency between experts and us for each entity is bolded.

| Entity Type | Inconsistent Number Due to Different Factors |  |  | Total | Experts' False Annotation Rate(#Experts False annotation/#Total,%) | Discrepant Rules Rate(#Discrepant Rules/#Total,%) |
| --- | --- | --- | --- | --- | --- | --- |
|  | Discrepant Rules | Experts' False Annotation | Our False Annotation |  |  |  |
| Gene | **14** | 2 | 8 | 24 | 8.33 | 58.33 |
| Variant | **26** | 1 | 0 | 27 | 3.70 | 96.30 |
| Species | **19** | 3 | 2 | 24 | 12.50 | 79.17 |
| Disease | **160** | 58 | 7 | 225 | 25.78 | 71.11 |
| Total | 219 | 64 | 17 | 300 | 21.33 | 73.33 |

### 2, The statistics of annotated entities in datasets at the finetuning stage

Table 3 shows the statistics of annotations in datasets for Phase I and Phase II of the finetuning stage, corresponding to ∼500,000 BERN-annotated PubMed abstracts and 400 our annotated full articles, respectively. It is obvious that the training, validation and test datasets in both phases at the finetuning stage are relatively balanced among the four entity types, respectively, which is essential for the current neural network (NN) models to achieve decent performance. In addition, the number of annotations in the training, validation and test datasets in Phase I of the finetuning stage is 30∼50 times, 25∼50 times and 20∼75 times of that in Phase II of the finetuning stage, respectively, indicating that the annotated corpus for the finetuning stage is considerably enriched from our small-size high-quality dataset by adding publicly available machine annotated corpus.

**Table 3.**
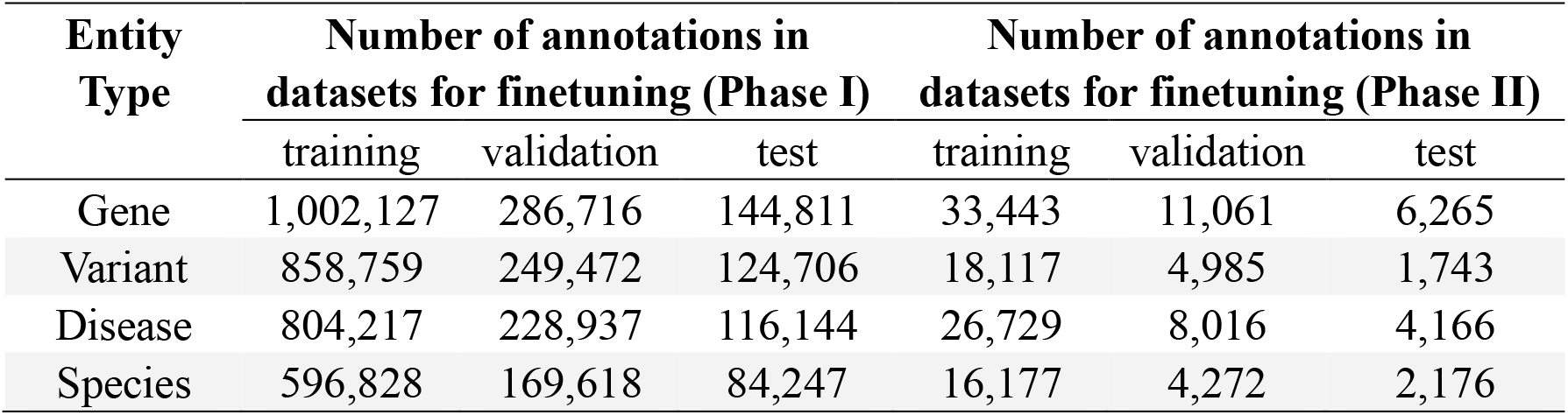
the statistics of annotations in datasets for Phase I and Phase II of the finetuning stage, corresponding to ∼500,000 BERN-annotated PubMed abstracts and 400 our annotated full articles, respectively.

### 3, Performance comparison of BERT-based NER, DistilBert-based NER and BERN

Table 4 exhibits the performance comparison of BERT-based NER, DistilBERT-based NER, and BERN^R^ for gene, variant, disease and species at the entity level. BERN^R^ indicates BERN reported metrics.^29^ It is clear that the F1-scores of BERT-based NER for gene, disease and species are all highest while the F1 score of BERT-based NER for variant is nearly 0.2% lower than that of BERN^R^. The F1-scores of NER of gene, variant, disease and species for the BERT-based model are 97.28%, 93.52%, 92.54% and 95.76%, respectively, while those for the DistilBERT-based model are 95.14%, 86.26%, 91.37% and 89.92%, respectively. Therefore, the F1 scores of the DistilBERT-based NER model retains 97.8%, 92.2%, 98.7% and 93.9% of those of BERT-based NER for gene, variant, disease and species, respectively.

**Table 4.** the performance comparison of BERT-based NER, DistilBERT-based NER, and BERN^R^ for gene, variant, disease and species at the entity level. BERN^R^ indicates BERN reported metrics.

| Entity Type | Model | Precision(%) | Recall(%) | F1-Score(%) |
| --- | --- | --- | --- | --- |
| Gene | BERT | **97.11** | **97.45** | **97.28** |
|  | DistilBERT | 94.84 | 95.45 | 95.14 |
|  | BERN<sup>R</sup> | 85.16 | 83.65 | 84.40 |
| Variant | BERT | 94.31 | **92.75** | 93.52 |
|  | DistilBERT | 85.94 | 86.59 | 86.26 |
|  | BERN<sup>R</sup> | **97.25** | 90.40 | **93.70** |
| Disease | BERT | **91.22** | **93.90** | **92.54** |
|  | DistilBERT | 90.44 | 92.32 | 91.37 |
|  | BERN<sup>R</sup> | 89.04 | 89.69 | 89.36 |
| Species | BERT | **98.30** | **93.34** | **95.76** |
|  | DistilBERT | 96.39 | 84.26 | 89.92 |
|  | BERN<sup>R</sup> | 93.84 | 86.11 | 89.81 |

## Discussion

Training deep learning NN models often requires tremendous resources and time. Fortunately, pretrained models based on huge corpora are often readily reused in NLP community. For instance, in order to adapt BERT models to the domain-specific language of biomedical texts, BioBERT was re-pretrained with PubMed abstracts and PubMed central full-text articles based on the BERT pretrained model.^28^ We started our model by loading pretrained BioBERT model weights. Although the overall model structure we selected is the same as BioBERT, the performance of our model is much better than that of BioBERT. As indicated in Table 4, the F1-scores of BERT-based NER for gene, disease and species are 12.88%, 3.18% and 5.95% higher than those of BioBERT in BERN^R^, respectively. Actually, BERN predictions on the same test set as in BERT-based NER were also obtained from BERN server. However, its performance was not as good as that of BERN^R^, so the corresponding metrics are not included here. It is noted that the F1-score for variant is not compared since BERN^R^ adopted the tool of tmVar 2.0 for variant recognition, although the F1-score for the BERT-based NER model— 93.52%— is comparable to that of BERN^R^— 93.70%.

There are at least two reasons for the above improvement. Firstly, we have carefully curated a high-quality data set to finetune the Bert-based NER and DistilBert-based NER models instead of using public available scattered entity-type corpora without the key entity of variant as in Biobert. The improvement indicates the significance of the good quality of training data for a NN-based model, especially for domain-specific corpora. Secondly, our model trained all four entity types jointly. An obvious advantage is that the features of all entities can be shared so that the internal relationships among the entities can be captured, which actually provides much more information than a separately-trained model. Another advantage is that the joint model can be easily further extended to relationship extraction tasks where at least two entity types should be included.

While the F1-scores of DistilBERT-based NER are relatively smaller than the corresponding values of its larger counterpart, the F1-scores of DistilBERT-based NER for gene, disease and species are still 10.74%, 2.01% and 0.11% higher than those of BioBERT in BERN^R^, respectively. This demonstrates the effectiveness of knowledge distillation of a large pretrained general-purpose language representation model, suggesting that the well-trained DistilBERT-based NER model can be applied for online inference.

Several directions for future work can be proposed based on the current study. Firstly, manually annotating is a time-consuming process, and thus automated algorithms can be introduced to accelerate the process. Secondly, the optimized Bert-based NER model may be applied to a large number of literatures to build a more accurate web-based platform for entity tagging for the biomedical community. Finally, the DistilBERT-based NER model can be added on the web-based platform for real-time, raw text tagging due to its faster computing speed.

## Conclusion

We report a combined high-quality manual annotation and deep-learning NLP study to make accurate NER for biomedical literatures. A total of 400 full articles from PubMed are annotated based on our home-made entity annotation guidelines. Both a BERT-based large model and a DistilBERT-based simplified model were constructed, trained and optimized for offline and online inference, respectively. The performance for both our BERT-based NER model and DistilBERT-based NER model outperforms that of the state-of-art model—BioBERT, indicating the significance to train an NER model on biomedical domain literatures jointly with high-quality annotated datasets. It is quite promising that the models can be applied to construct a useful and efficient entity-tagging platform.

## Acknowledgements

This work was supported by the National Natural Science Foundation of China (32001076). We would like to thank our colleague Haiping Huang for insightful discussion.

## Author contributions

D.L. Huang conceived, conducted the NER experiments and wrote the manuscript. Q.L. Zeng, Y. Xiong, S.X. Liu, C.Q. Pang, M.L. Xia, T. Fang, Y.L. Ma, C.C. Qiang, Y. Zhang, Y. Zhang and H. Li constructed the annotation guidelines and annotated our corpus. D.L. Huang, Q.L. Zeng and C.Q. Pang analyzed the results. All authors reviewed the manuscript.

## References

1 Lappalainen, T., Scott, A. J., Brandt, M. & Hall, I. M. Genomic analysis in the age of human genome sequencing. Cell 177, 70–84 (2019).

2 Good, B. M., Ainscough, B. J., McMichael, J. F., Su, A. I. & Griffith, O. L. Organizing knowledge to enable personalization of medicine in cancer. Genome biol. 15, 1–9 (2014).

3 Richards, S. et al. Standards and guidelines for the interpretation of sequence variants: a joint consensus recommendation of the American College of Medical Genetics and Genomics and the Association for Molecular Pathology. Genet. Med. 17, 405–423 (2015).

4 Allot, A. et al. LitVar: a semantic search engine for linking genomic variant data in PubMed and PMC. Nucleic Acids Res. 46, W530–W536 (2018).

5 den Dunnen, J. T. et al. HGVS recommendations for the description of sequence variants: 2016 update. Hum. Mutat. 37, 564–569 (2016).

6 Landrum, M. J. et al. ClinVar: public archive of interpretations of clinically relevant variants. Nucleic Acids Res. 44, D862–D868 (2016).

7 Li, Q. & Wang, K. InterVar: clinical interpretation of genetic variants by the 2015 ACMG-AMP guidelines. Am. J. Hum. Genet. 100, 267–280 (2017).

8 Ahern, C. & Brokamp, E. The Utility of Genomic Variant Databases in Genetic Counseling. (2016).

9 Bean, L. J. & Hegde, M. R. Gene variant databases and sharing: creating a global genomic variant database for personalized medicine. Hum. Mutat. 37, 559–563 (2016).

10 Goulart, R. R. V., de Lima, V. L. S. & Xavier, C. C. A systematic review of named entity recognition in biomedical texts. J. Brazilian Comput. Soc. 17, 103–116 (2011).

11 Wang, X. et al. Cross-type biomedical named entity recognition with deep multi-task learning. Bioinformatics 35, 1745–1752 (2019).

12 Xu, K., Yang, Z., Kang, P., Wang, Q. & Liu, W. Document-level attention-based BiLSTM-CRF incorporating disease dictionary for disease named entity recognition. Comput. Biol. Med. 108, 122–132 (2019).

13 Sachan, D. S., Xie, P., Sachan, M. & Xing, E. P. Effective use of bidirectional language modeling for transfer learning in biomedical named entity recognition. Proceedings of Machine Learning Research 85, 1–19 (2018).

14 Colic, N., Furrer, L. & Rinaldi, F. Annotating the Pandemic: Named Entity Recognition and Normalisation in COVID-19 Literature. (2020).

15 Kim, J.-D., Ohta, T., Tsuruoka, Y., Tateisi, Y. & Collier, N. in Proceedings of the international joint workshop on natural language processing in biomedicine and its applications. 70–75 (Citeseer).

16 Song, M., Yu, H. & Han, W.-S. Developing a hybrid dictionary-based bio-entity recognition technique. BMC Med. Inf. Decis. Making 15, 1–8 (2015).

17 Song, H.-J., Jo, B.-C., Park, C.-Y., Kim, J.-D. & Kim, Y.-S. Comparison of named entity recognition methodologies in biomedical documents. Biomedical engineering online 17, 1–14 (2018).

18 Yadav, V. & Bethard, S. A survey on recent advances in named entity recognition from deep learning models. arXiv preprint arXiv:1910.11470 (2019).

19 Wei, C.-H., Allot, A., Leaman, R. & Lu, Z. PubTator central: automated concept annotation for biomedical full text articles. Nucleic Acids Res. 47, W587–W593 (2019).

20 Wei, C.-H., Kao, H.-Y. & Lu, Z. GNormPlus: an integrative approach for tagging genes, gene families, and protein domains. BioMed Res. Int. 2015 (2015).

21 Wei, C.-H. et al. tmVar 2.0: integrating genomic variant information from literature with dbSNP and ClinVar for precision medicine. Bioinformatics 34, 80–87 (2018).

22 Wei, C.-H., Kao, H.-Y. & Lu, Z. PubTator: a web-based text mining tool for assisting biocuration. Nucleic Acids Res. 41, W518–W522 (2013).

23 Chen, Q. et al. BioConceptVec: creating and evaluating literature-based biomedical concept embeddings on a large scale. PLoS Comp. Biol. 16, e1007617 (2020).

24 Chiu, J. P. & Nichols, E. Named entity recognition with bidirectional LSTM-CNNs. Transactions of the Association for Computational Linguistics 4, 357–370 (2016).

25 Schuster, M. & Paliwal, K. K. Bidirectional recurrent neural networks. IEEE transactions on Signal Processing 45, 2673–2681 (1997).

26 Cho, H. & Lee, H. Biomedical named entity recognition using deep neural networks with contextual information. BMC Bioinformatics 20, 1–11 (2019).

27 Devlin, J., Chang, M.-W., Lee, K. & Toutanova, K. Bert: Pre-training of deep bidirectional transformers for language understanding. arXiv preprint arXiv:1810.04805 (2018).

28 Lee, J. et al. BioBERT: a pre-trained biomedical language representation model for biomedical text mining. Bioinformatics 36, 1234–1240 (2020).

29 Kim, D. et al. A neural named entity recognition and multi-type normalization tool for biomedical text mining. IEEE Access 7, 73729–73740 (2019).

30 Hinton, G., Vinyals, O. & Dean, J. Distilling the knowledge in a neural network. arXiv preprint 1503.02531 (2015).

31 Sun, S., Cheng, Y., Gan, Z. & Liu, J. Patient knowledge distillation for bert model compression. arXiv preprint 1908.09355 (2019).

32 Sanh, V., Debut, L., Chaumond, J. & Wolf, T. DistilBERT, a distilled version of BERT: smaller, faster, cheaper and lighter. arXiv preprint 1910.01108 (2019).

33 Jiao, X. et al. Tinybert: Distilling bert for natural language understanding. arXiv preprint 1909.10351 (2019).

34 Li, J. et al. BERT-EMD: Many-to-Many Layer Mapping for BERT Compression with Earth Mover’s Distance. arXiv preprint 2010.06133 (2020).

35 Lee, K. et al. BRONCO: Biomedical entity Relation ONcology COrpus for extracting gene-variant-disease-drug relations. Database 2016 (2016).

36 Doğan, R. I., Leaman, R. & Lu, Z. NCBI disease corpus: a resource for disease name recognition and concept normalization. J. Biomed. Inf. 47, 1–10 (2014).

37 Gerner, M., Nenadic, G. & Bergman, C. M. LINNAEUS: a species name identification system for biomedical literature. BMC Bioinform. 11, 1–17 (2010).

38 Morgan, A. A. et al. Overview of BioCreative II gene normalization. Genome biol. 9, 1–19 (2008).

39 Buchholz, S. & Marsi, E. in Proceedings of the tenth conference on computational natural language learning (CoNLL-X). 149–164.

